# Patterns of shared signatures of recent positive selection across human populations

**DOI:** 10.1101/109371

**Authors:** Kelsey Elizabeth Johnson, Benjamin F. Voight

## Abstract

Scans for positive selection in human populations have identified hundreds of sites across the genome with evidence of recent adaptation. These signatures often overlap across populations, but the question of how often these overlaps represent a single ancestral event remains unresolved. If a single positive selection event spread across many populations, the same sweeping haplotype should appear in each population and the selective pressure could be common across diverse populations and environments. Identifying such shared selective events would be of fundamental interest, pointing to genomic loci and human traits important in recent history across the globe. Additionally, genomic annotations that recently became available could help attach these signatures to a potential gene and molecular phenotype that may have been selected across multiple populations. We performed a scan for positive selection using the integrated haplotype score on 20 populations, and compared sweeping haplotypes using the haplotype-clustering capability of fastPHASE to create a catalog of shared and unshared overlapping selective sweeps in these populations. Using additional genomic annotations, we connect these multi-population sweep overlaps with potential biological mechanisms at several loci, including potential new sites of adaptive introgression, the glycophorin locus associated with malarial resistance, and the alcohol dehydrogenase cluster associated with alcohol dependency.

## INTRODUCTION

Positive selection is the process whereby a genetic variant rapidly increases in frequency in a population due to the fitness advantage of one allele over the other. Recent positive selection has been a driving force in human evolution, and studies of loci targeted by positive selection have uncovered phenotypes that may have been adaptive in recent human evolutionary history (*e.g.*, [1–3]). One observation that has emerged from scans for positively selected loci that have not yet reached fixation [4–12] is that these signatures are often found across multiple populations, localized to discrete locations in the genome [6,7,10,11]. Large-scale sequencing data now available from diverse human populations offers the opportunity to characterize the frequency with which such overlapping signatures share a common, ancestral event - and potentially a common selective pressure. Identifying shared selective events would be of fundamental interest, pointing to genomic loci and human traits important in recent history across the globe. In addition, these selective targets could help to clarify the range of population genetic models compatible with observed data, in order to elucidate the demographic and selective forces that shape global genomic diversity.

Recently generated annotations of the human genome also offer the added potential to identify candidate genes or variants targeted by selection and their associated mechanism. For example, the influx of genomic data on expression quantitative trait loci (eQTLs) across many tissue types [13], and/or inferred regions of ancient hominin introgression [14–20] now provide a richer foundation to investigate the potential biological targets under selection at these loci. While identifying the causal variant at a site of positive selection is notoriously difficult, if SNPs on a selected haplotype are associated with changes in expression of a nearby gene, this information could help attach the signature to a potential gene and molecular phenotype. Small insertions and deletions (indels), and copy number variants on sweeping haplotypes also represent potentially functional variation that could be targeted by natural selection.

In this study, we focus on the detection of genomic signatures compatible with selection on a newly introduced mutation in humans that has not yet reached fixation (*i.e.*, hard, ongoing sweeps) to explore their distribution across populations and spanning the genome. We performed a scan for positive selection using the integrated haplotype score (iHS) on 20 populations from four continental groups from Phase 3 of the 1000 Genomes Project (1KG) [21]. We found that 88% of sweep events overlapped across two or more populations, correlating with population relatedness and geographic proximity. 59% of overlaps were shared (i.e., a similar sweeping haplotype was present) across populations, and 29% of overlaps were shared across continents. Using additional genomic annotations, we connect these multi-population sweep overlaps with potential mechanisms at (i) the glycophorin cluster (*GYPA*, *GYPB*, and *GYPE*), where we observe sweeps across all four continental groups in a region associated with malarial resistance; (ii) sweeps across African populations at the X chromosome gene *DGKK*, implicated in the genital deformity hypospadias in males; (iii) a sweep shared in European populations tagged by a coding variant in the gene *MTHFR*, which is associated with homocysteine levels and a multitude of additional traits; (iv) two putative regions of adaptive introgression from Neandertals; and (v) the alcohol dehydrogenase (*ADH*) cluster, where a sweep in Africa is associated with alcohol dependence in African Americans.

## RESULTS

### A catalog of signals of recent positive selection across human populations

To identify genomic intervals with extended haplotypes compatible with the action of recent, positive selection, we measured iHS normalized separately for the autosomes and X chromosome across 26 populations from the 1KG project (**Methods**, **S1 Table**). We normalized iHS scores with an added correction for local recombination rate, in response to an observed excess of extreme iHS scores at regions of low local recombination rate (**Methods**). For each population’s iHS scan, we identified putative sweep intervals that segregated an unusual aggregation of extreme values of iHS (**Methods**, **S3 Table)**. Consistent with previous reports [4], the number of sweep intervals per population correlated with its effective population size (**S4 Table**). We defined the tag SNP for each interval as the highest scoring variant by the absolute value of iHS, as we expect the tag SNPs to be in strong linkage disequilibrium (LD) with the putative causal, selected variant of the sweep. Our sweep intervals recovered 11 of the 12 top signatures reported in the original iHS paper [4], and 14 of the top 22 signatures reported in [22]. We observed more extreme iHS scores using WGS data compared to array-based genotype data. For example, previously in CEU only 6 of 256 signatures (2.3%) had an absolute value of iHS > 5 for their most extreme score [4], while in our scan 92 of 597 signatures (15%) had an absolute value > 5 for their most extreme score, and 28 had a most extreme iHS score with absolute value > 6.

### Signatures of recent positive selection frequently overlap within continental groups

We next sought to characterize the frequency that putative sweep intervals overlap the same genomic region across multiple populations. Here, we excluded recently admixed populations (ASW, ACB, MXL, PUR, CLM) as events observed in those groups could simply reflect selection in ancestral populations predating admixture (**Methods**). We found that related populations (as measured by F_ST_) more often overlap in their putative sweeps intervals, relative to more distantly related pairs, an observation consistent with previous findings [6,7,10] (Fig 1). To explain the residual variability in sweep overlaps, we performed multiple linear regression to model the fraction of sweeps overlapping between all pairs of populations, using pairwise F_ST_, continental grouping (*e.g.*, within East Asia, or between Europe and Africa, etc.), straight-line geographic distance, difference in latitude, and difference in longitude as potential predictors for the fraction of sweep intervals that overlap. The most significant predictor was the continental-grouping label (P < 2 × 10^−16^), along with pairwise F_ST_, difference in latitude, and difference in longitude explaining additional variance in the fraction of sweep intervals that overlap. A final model with these four variables explained virtually all variability in the fraction of overlaps (R^2^ = 0.96, P < 2.2 × 10^−16^, Fig 1).

**Fig 1.**
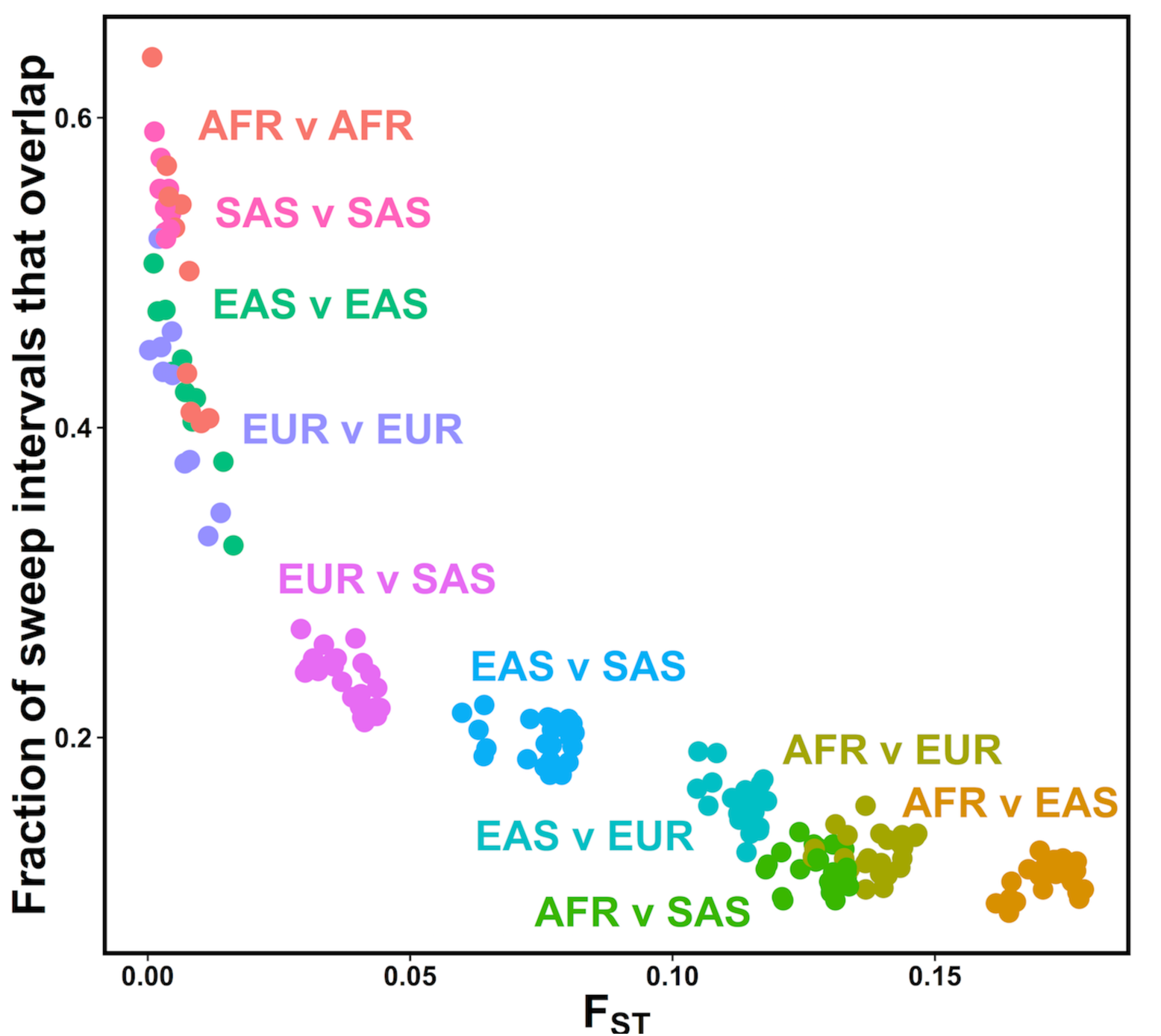
Closely related populations have sweep overlaps more frequently. For each population pair, the fraction of sweep intervals that overlap is plotted against pairwise estimated F_ST_. Each population pair (dots) are colored by their continental groupings (e.g. EUR v SAS = one European population *vs*. one South Asian population).

### Overlapping sweep intervals across human populations occur in genomic hotspots

In the process of characterizing sweep overlaps across the genome, we observed cases where many sweeps appeared to cluster in specific genomic locations. The most striking clustering of overlapping sweep intervals occurred on chromosome 17 (Fig 2A), with 23 overlapping events in total, of which 14 span continental groups. While previous reports have investigated how often sweeps overlap across the globe, the extent to which putative sweep intervals are organized and/or cluster across the genome has not been previously quantified. To model this phenomenon, we measured the rates at which individual or overlapping sweep intervals occurred across the genome, fitting the observed distribution of the number of events in 10Mb windows with individual or mixtures of Poisson distributions (**Methods**). As a positive control, we first modeled the count of genes in each window. We found that a mixture model with five components best fit the frequency at which genes occur in the genome (**Methods**), an expected result owing to the fact that genes are indeed not uniformly distributed across the genome. Turning next to the frequency of sweeps in our genome, we first observed that the counts of single population sweep intervals in a window were best modeled by a single rate genome-wide (Fig 2B). In contrast, sweeps overlapping across populations were best fit by a mixture of Poisson distributions with three different rates (P = 2.7 × 10^−7^, χ^2^ test vs. two component mixture, Fig 2C, **Methods**). These overlap hotspots were not explained by the number of genes in a window (Pearson’s correlation = 0.07, P = 0.22). These data indicate that putative sweep intervals overlapping multiple human populations appear to aggregate in discrete “hotspots” of activity.

**Fig 2.**
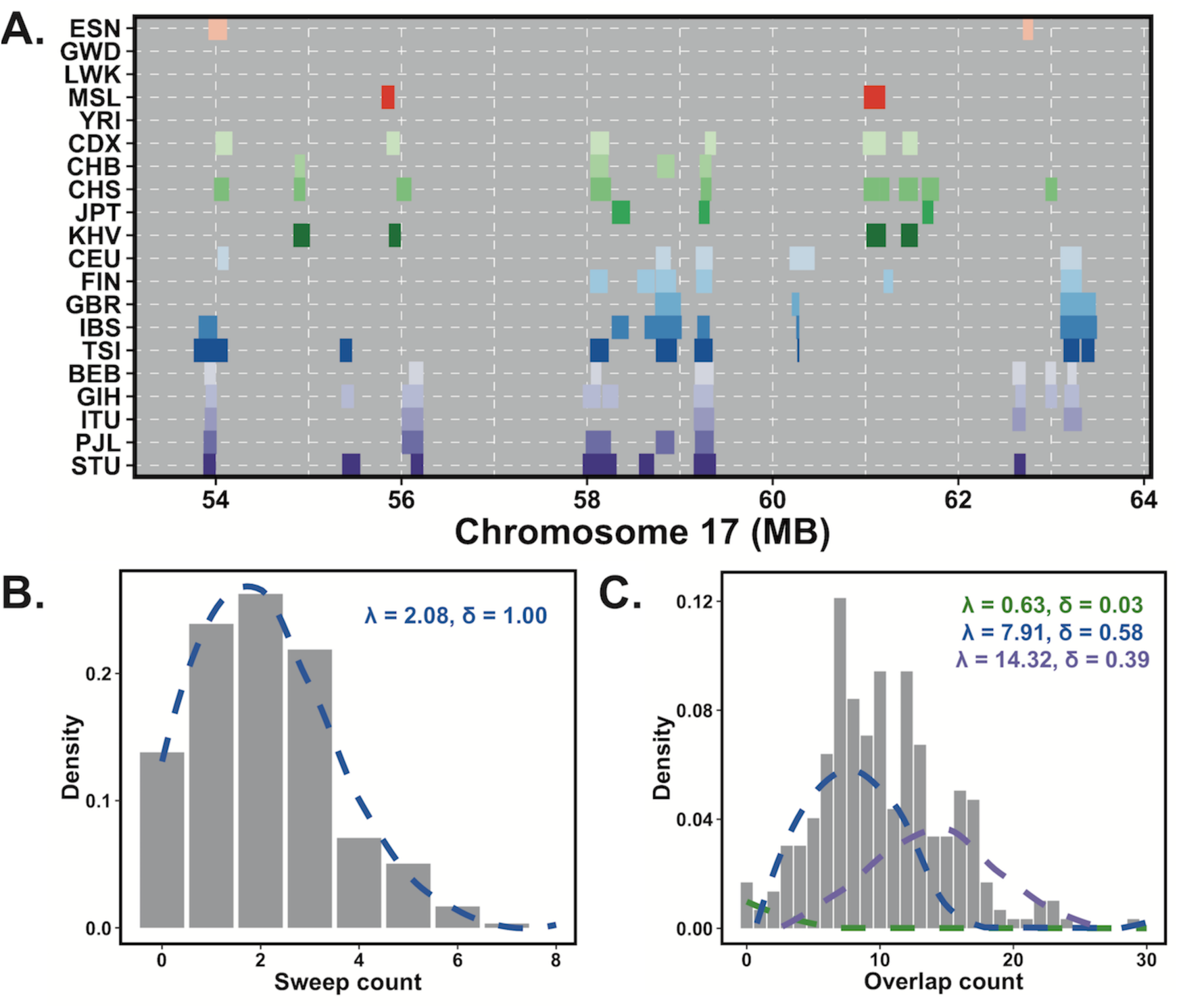
Overlapping sweeps tend to cluster in the genome. **(A)** An example of a 10 megabase (Mb) window on chromosome 17 with multiple overlaps across many populations. **(B)** The distribution of sweep interval counts in 10 Mb windows across the genome for a single population (LWK). The histogram plots the observed counts, and the blue dashed line is the best-fit Poisson distribution. **(C)** The distribution of sweep overlaps across two or more populations in 10 Mb windows across the genome. The histogram plots the observed counts, and the dashed lines represent the results of Poisson mixture modeling. The best-fit model was the three-component model shown here.

### Complex patterns of sweep sharing across populations and continents

We next sought to identify selective sweeps that are potentially shared across populations, *i.e.*, where the putative sweeping haplotype is similar across populations. Sharing could occur in several ways, including a common ancestral event occurring before population divergence that persisted to the present day, or via gene flow of advantageous alleles between populations. To characterize haplotype similarity across populations at our genomic intervals tagged by unusual iHS scores, we utilized the program fastPHASE [23]. Using a hidden Markov model, fastPHASE models the observed distribution of haplotypes as mosaics of *K* ancestral haplotypes, which allows us to map the SNP that tags the sweep interval to an ancestral haplotype jointly across multiple populations at once without arbitrarily choosing a physical span to build a tree of haplotypes or otherwise measure relatedness **(Methods)**.

Overall, out of 1,803 intervals shared across populations, 521 (29%) were shared across continents, most frequently between Europe and South Asia and consistent with observed lower genetic differentiation relative to other continental comparisons (**S5 Table)**. Indeed, consistent with our previous analysis using all intervals, the fraction of sweep overlaps that were shared between a pair of populations was strongly correlated with F_ST_ (S1 Fig). To determine if the observed extent of sweep sharing was unusual, we applied our fastPHASE haplotype labeling procedure to random sites across the genome for each population pair, matched for distance to gene, interval size, tag SNP frequency, and derived/ancestral allele distribution as the observed sweep overlaps. For all intra-continental population pairs, and all but one Eurasian inter-continental pair, the degree of sweep sharing was higher than the background rate (Fig 3, **S6 Table**), suggesting that the sweep sharing we observe is not driven purely by haplotype similarities across closely related populations.

**Fig 3.**
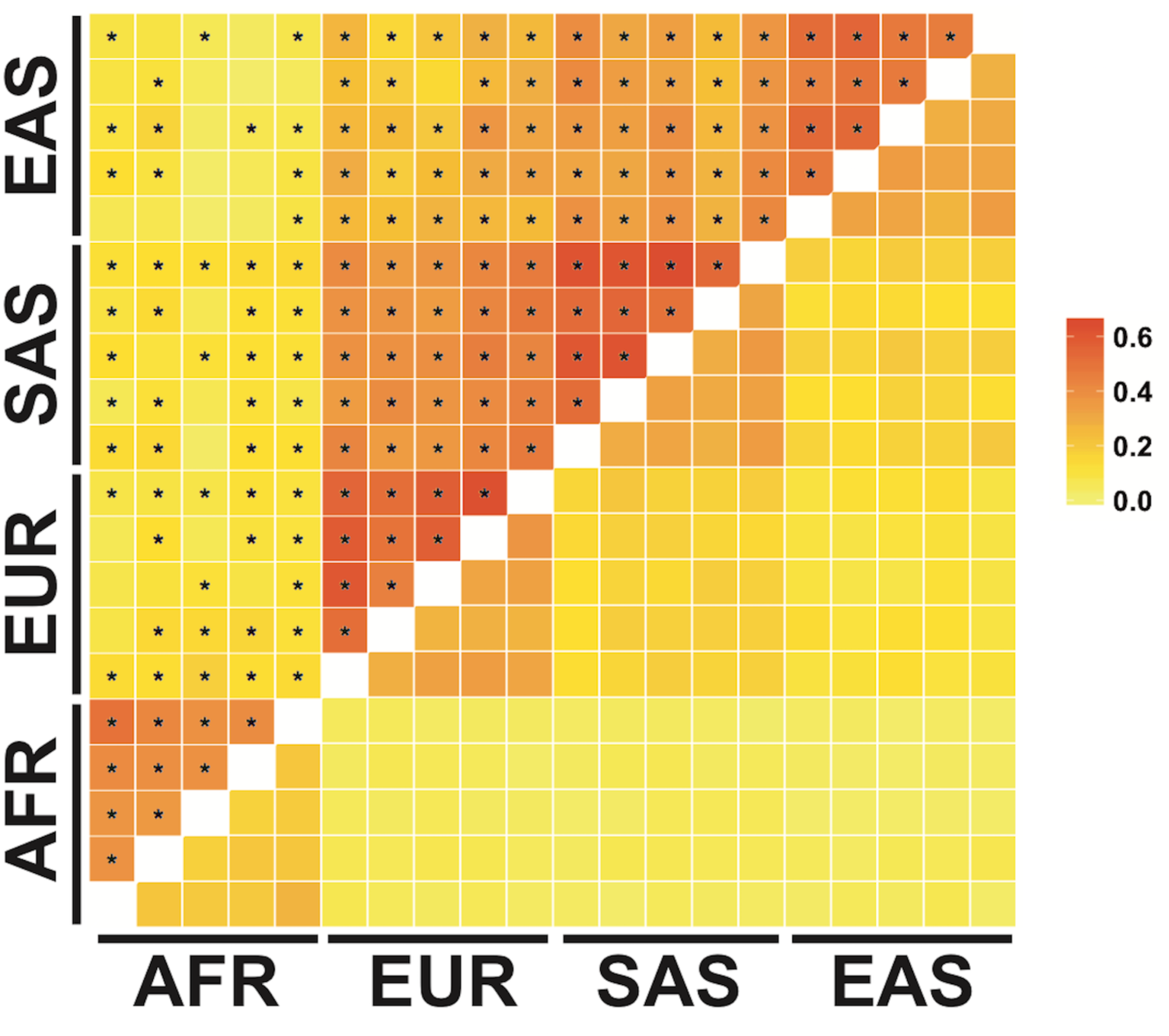
Enrichment of shared sweeps across population pairs. Squares below the diagonal represent the null fraction of overlaps shared across population pairs, from randomly placed overlaps across the genome. Squares above the diagonal represent the observed fraction of sweep overlaps shared for each population pairs. Squares are marked with an asterisk if the observed fraction shared was significantly higher than the null distribution. Populations are arranged alphabetically top to bottom, right to left within continental groups.

Though the majority of inter-continental shared sweeps are across non-African populations, we did observe examples of shared sweeps between African and non-African populations. In total, 9.4% of observed sweep overlaps between African and non-African population pairs were called as shared (491 total), compared with 4.0% of control overlaps (99% CI: 3.7-4.4%). For example, on chromosome 1 at ~47MB, a sweeping haplotype shared across African and European populations fell in a cytochrome P450 gene cluster, including *CYP4B1*, *CYP4Z2P*, *CYP4A11*, *CYP4X1*, *CYP4Z1*, and *CYP4A22* (S2 Fig). Selection at this site in Europeans, and an enrichment of unusual iHS signatures in the cytochrome P450 gene family, has been previously described [4]. The SNPs shared across the sweeping haplotypes are also eQTLs for *CYP4X1*, *CYP4Z1,* and *CYP4A22-AS1*, with strong LD (R^2^ > 0.8) between the populations’ tag iHS SNPs and the lead eQTLs for *CYP4X1* and *CYP4A22-AS1* in testis (S2 Fig). With a catalog of shared and overlapping selective sweeps in hand, we next aimed to identify specific regions of sweep sharing that connected the interval to a gene, pathway, or phenotype when considered alongside annotations of the genome (*e.g.*, gene expression, complex-trait phenotype associations, sequences of Neandertal introgression, or pathway enrichment.).

### Shared and overlapping sweeps in a region implicated in malarial resistance

With a sweep overlap across thirteen populations from all four continental groups, the glycophorin gene cluster (*GYPA/GYPB/GYPE*) on chromosome 4 came to our attention for its repeated targeting by positive selection and its prior implication in malaria resistance (Fig 4). This genomic region has been noted as a target of positive selection in humans [24–27], and as a target of ancient balancing selection shared between humans and chimpanzees [28]. In our study, the sweep in IBS and South Asians was on a shared haplotype, while the African populations, CHB, and CEU had unique sweeping haplotypes (Fig 4). The sweeping haplotypes from all four continental groups carried eQTLs for *GYPB* and *GYPE*. The sweeping haplotypes present in these populations also carried several nonsynonymous mutations in GYPB; however, these mostly occurred at low frequency (<2%) and thus were not likely to be the selected causal variant. The one exception, rs7683365, has a minor allele frequency in these selected populations ranging from 4% in CHB to 38% in ITU. rs7683365 specifies S>s antigen status and has been found to be associated with susceptibility to malaria infection in an admixed Brazilian population [29]. This variant is in nearly complete LD with the tag variant (D’ > 0.97) in CHB, CEU, and IBS, one African populations (MSL), and three South Asian populations (GIH, ITU, PJL).

**Fig 4.**
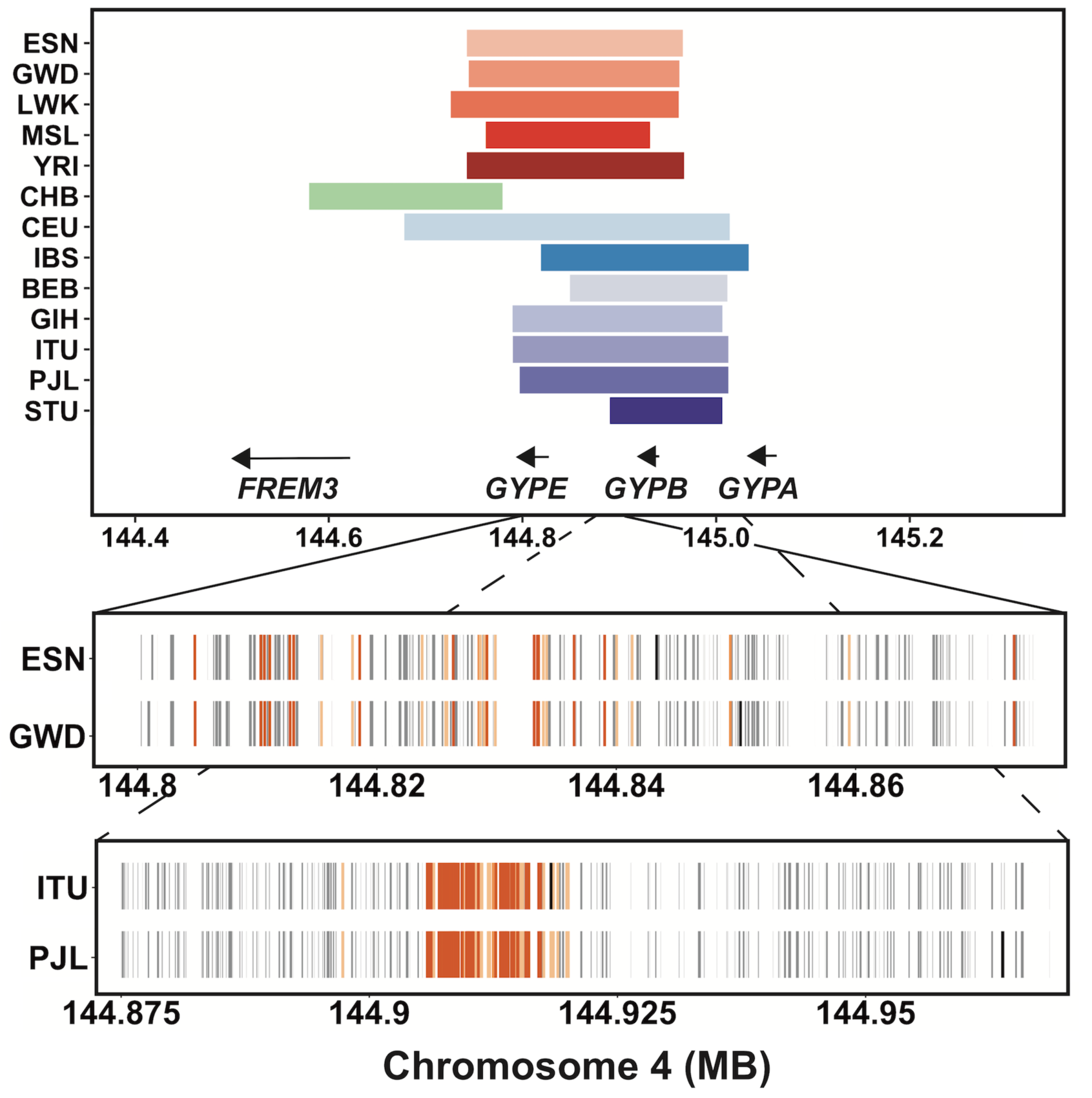
Signatures of positive selection at the *GYP* locus on chromosome 4. We observed signatures of positive selection in 13 populations at the GYP locus, including at least one population from each studied continental group. The top panel shows the sweep intervals of populations with sweeps at this locus, and the positions of the *GYP* genes. The bottom panels show the sweeping haplotypes for two African (ESN, GWD) and two South Asian (ITU, PJL) populations’ within-continent shared sweeps. The gray tick marks in each populations’ row indicate the presence of a derived allele on the sweeping haplotype most common in that population, with a black tick indicating the position of each population’s SNP with the most extreme iHS value. Also shown in orange are the significant eQTLs for *GYPE* (light orange) or both *GYPB* and *GYPE* (dark orange) in LD with these population’s shared haplotype (D’ = 1). The eQTLs for *GYPB* and *GYPE* are from whole blood.

A GWAS of malaria phenotypes found a significant association located between *FREM3* and *GYPE* in sub-Saharan Africans [30], tagging a duplication event that also appears to have undergone recent positive selection in Kenyans [27]. A reference panel with diverse African representation was sequenced, with copy number inferences in strong agreement with the CNV calls from 1KG [27]. However, we found that the CNVs present in 1KG were rare (frequency < 5%) or not in strong LD with the sweep tag variants (R^2^ < 0.7), suggesting that the copy number variants present in the 1KG dataset were not likely to be the causal variants at these iHS signatures. In sum, we identified multiple signatures of positive selection on distinct haplotypes in all four continental groups (**S5 Table**), at most of which the causal variants did not appear to be coding or structural variants. These sweeping haplotypes all carry eQTLs for *GYPB* and *GYPE*, genes previously implicated in association to malarial resistance and ancient balancing selection.

### Intersection of signatures of positive selection with the GWAS catalog

Genetic variants associated with disease represent an additional resource for interpretation of signatures of positive selection. Previous work has indicated an enrichment of extreme iHS scores at GWAS signals for autoimmune diseases [31], and we hypothesized that this or other traits might be enriched for GWAS signatures linked to our signatures of positive selection. In total, 186 sweep tag SNPs from all 20 populations (out of 11,655; 1.6%) were in strong LD with at least one genome-wide significant GWAS SNP (R^2^ ≥ 0.9, **S8 Table**). None of the traits we investigated showed clear, compelling evidence of enrichment (**Methods)**. However, this intersection did identify specific candidates for the potential phenotype of selection at those loci. In one example, a sweep overlap across all five African populations falls at the gene diacylglycerol kinase kappa (*DGKK*) on the X chromosome (S3 Fig). Variants in this gene have been associated in Europeans with hypospadias [32,33], a prevalent birth defect of ectopic positioning of the opening of the urethra in males. A lead SNP for the hypospadias association at *DGKK*, rs4554617, is in perfect LD (R^2^ =1) with the sweep tag SNP in YRI at this overlap. The unselected, ancestral allele (C) is associated with increased risk for hypospadias in Europeans (OR = 2.52, P = 1.01×10^−93^) [33]. The sweeping haplotypes in this overlap were classified as shared for MSL and LWK, and unique in ESN, GWD, and YRI; and this variant is only in strong LD with the tag SNP for YRI. The sweeping haplotypes for these genes do not carry any coding variants; and while they contain eQTLs for several other genes in the region, we did not find evidence for eQTLs for *DGKK* (**Methods**). Though multiple independent studies have found an association with hypospadias at this locus, the pathogenic mechanism remains unknown.

A second example occurred at the methylenetetrahydrofolate reductase (*MTHFR*) gene on chromosome 1, where a nonsynonymous variant (A222V, rs1801133) has been extensively studied for its association with homocysteine levels [34,35]. A sweep overlap at this locus with three European populations (CEU, GBR, IBS) and JPT was called as shared across all four populations (S4 Fig). rs1801133 is the tag SNP for CEU and GBR’s sweeps, and is in moderate LD with IBS’ tag SNP (R^2^=0.41, D’=1). Though a sweep interval was not called for TSI at this locus, it also has an unusually extreme iHS score for rs1801133 (iHS = -3.311). *MTHFR* encodes an enzyme involved in folate metabolism, and the derived allele that appears to be under selection in Europeans (T) is associated with higher homocysteine levels [34,35], lower folate vitamin and B12 levels [36,37], and multiple additional traits (**Discussion**).

### Evidence for adaptive introgression from Neandertals in non-African populations

Gene flow occurred between humans and other hominins after migration out of Africa, resulting in ~2% of non-African humans’ genomes deriving from Neanderthal or Denisovan origin [14–20]. Examples of positive selection on introgressed genetic variation have shown that positive selection acted on genetic variation from ancient hominins at some loci [38–43]. While some of these examples are confined to a single population (e.g. *EPAS1* in Tibetans [43]), most are common across multiple populations [44], and thus we hypothesized that a subset of our shared sweeps could be additional examples of adaptive introgression. Using introgressed haplotypes from Neanderthals inferred in individuals from phase 3 of the 1000 Genomes Project (less than 1% of introgressed sequences in these populations are predicted to be of Denisovan origin [19]), we identified 141 candidate sweeps in LD with introgressed haplotypes (R^2^ ≥ 0.6, excluding X chromosome, **Methods, S9 Table**). These introgressed haplotypes include previously described adaptive targets such as the *HYAL2* locus on chromosome 3 in East Asians [41] and *OAS1* in Europeans [39]. We did not observe an overall enrichment of these introgressed haplotypes in our iHS intervals (P = 0.59, **Methods**), suggesting that introgression alone did not increase the likelihood of a haplotype to be identified as an unusual iHS signature. Of these 141 loci, we illustrate two candidate sweeps that had LD between an introgressed haplotype and a shared sweep across the most populations, along with functional information that generated a hypothesis for the target gene, variant, or phenotype. The first example occurred on chromosome 3, near cancer/testis antigen 64 (*CT64*, S5 Fig). At this locus a shared sweep between Europeans and South Asians tagged an adjacent introgressed Neandertal haplotype (R^2^ between 0.8 - 1.0). The sweep tag SNPs are also in strong linkage with eQTLs for *CT64*, a non-coding RNA primarily expressed in the testes. We observed strong concordance between the frequencies of the introgressed haplotype, sweep tag SNPs, and lead eQTL SNP in European populations (ranging from ~25-30%), and less strong concordance in the South Asian populations, where the sweep and introgressed haplotypes are at lower frequencies (~6-15%). This evidence suggests a variant introgressed from Neandertals that may have regulatory potential for *CT64* underwent a selective sweep in Europeans, and perhaps a less strong sweep in South Asians. The second example was found at ~41MB on chromosome 1, where all five South Asian populations have evidence of an introgressed haplotype at low to moderate frequency (ranging from 18% in BEB, GIH, ITU, STU to 30% in PJL). A sweeping haplotype intersecting this introgression event is also shared across all five populations (S6 Fig). Three introgressed SNPs on this haplotype (rs11209368, rs2300659, rs10493094) are nominally associated with chronic obstructive asthma and chronic airway obstruction [45] and are tightly linked (R^2^ ranging from 0.77 to 1.0) to the tags for the sweeping haplotypes in each of the five South Asian populations. The strongest association was the Neandertal allele at rs2300659, which had a p-value of 9×10^−4^ for chronic airway obstruction (OR = 1.24), and p-value of 3.02×10^−3^ for chronic obstructive asthma (OR = 1.55). Their shared haplotype also carries eQTLs for five genes: CTPS1, FOXO6, RP11-399E6.1, SCMH1, and SLFNL1. The eQTLs on the sweeping haplotype fall within several predicted regulatory elements in the region, including some with histone modifications in primary lung tissue [46].

### Overlapping and shared sweeps enriched in the ethanol oxidation pathway

We next sought to explore possible biological pathways targeted by shared selective events. As a large fraction of causal variants under positive selection are potentially non-coding [5,8,47], we hypothesized that regulatory variation in the form of eQTLs could indicate a potential causal, functional variant and/or gene target. We identified genes with cis-eQTLs from all tissue types in the GTEx dataset that were linked with shared sweeps (R^2^ ≥ 0.9) and tested for overrepresentation of biological pathways in this set of genes using ConsensusPathDB [48].

Excluding the human leukocyte antigen (HLA) genes (**Methods**), the most significant pathway was ethanol oxidation (P = 2.0 × 10^−5^, q-value = 0.047). This pathway includes six members of the alcohol dehydrogenase (*ADH*) gene cluster (1A, 1B, 1C, 4, 6, 7), as well as distinct genomic locations which include two aldehyde dehydrogenases (*ALDH2*, *ALDH1A1*) and two acyl-CoA synthetase short-chain family members (*ACSS1* and *ACSS2*). Of these 10 genes, 7 were included in our shared sweeps gene set (*ADH1A, ADH1C, ADH4, ADH6, ALDH2, ACSS1, ACSS2*). We also observed shared sweeps linked (D’ > 0.99) with eQTLs for *CYP2E1*, a primary enzyme in an alternative alcohol metabolism pathway. Moreover, *ADH1B*, *ADH4*, *ADH5*, and *ADH6* segregate eQTLs that intersect sweeping haplotypes that are unique to one population. The ADH gene cluster contains a previously described East Asian selective event targeting rs1229984 [49], a nonsynonymous variant in *ADH1B* (or alternatively, the noncoding *ADH1B* promoter variant rs3811801 [50]). A recent report also found evidence for an independent selective event for rs1229984 in Europeans [51]. In this cluster, we observed independent sweeping haplotypes in YRI and ESN at *ADH4* and *ADH5*, and a second distinct sweeping haplotype in YRI overlapping *ADH1B*, *ADH1C*, and *ADH7* (Fig 5).

**Fig 5.**
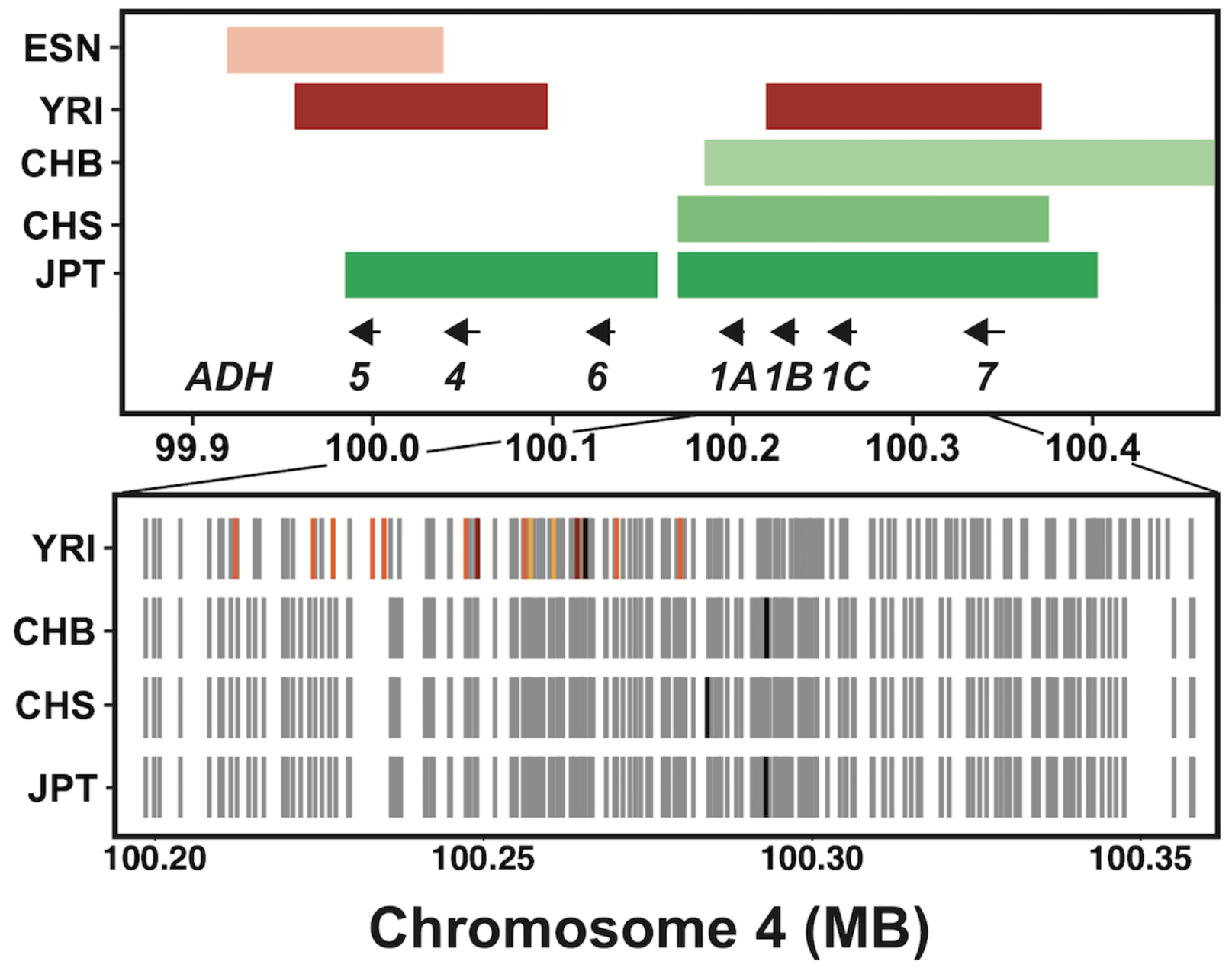
Signatures of positive selection at the *ADH* locus on chromosome 4. The top panel shows the sweep intervals of populations with sweeps at this locus, and the positions of the seven *ADH* cluster genes. The bottom panel shows the sweeping haplotypes of the four populations with sweeps in this region, with grey tick marks indicating the derived alleles present on the most common sweeping haplotype in that population. The black tick marks indicate the position of the SNP with the most extreme iHS score in each population. For YRI, the positions of significant *ADH4* and *ADH1C* eQTLs in subcutaneous adipose tissue (light orange), GWAS SNPs from Gelernter et al., 2014 (dark orange), and SNPs that are eQTLs for both genes and are GWAS SNPs (red) in LD with YRI’s tag SNP (R^2^ > 0.9) are shown.

As genome-wide association studies have identified genetic variation in the *ADH* locus associated with alcohol dependence (AD) [52–54], we next tested whether these associations were linked to the sweeping haplotype in YRI. The derived allele rs1229984-T of *ADH1B* is associated with increased *ADH1B* enzyme activity [55] and decreased risk of AD in East Asians [53,56]. In the YRI sweep interval spanning *ADH1B*, *ADH1C*, and *ADH7*, the leading iHS SNP (rs12639833, iHS = -5.133) was significantly associated with decreased risk for AD in African Americans [54], and we noted that the derived allele putatively under selection in YRI (rs12639833-T) also protected against AD. This association was specific to African Americans (though we note that this study also included European Americans [54]). rs12639833 lies in an intron of *ADH1C*, and is a significant eQTL for *ADH1C* (esophagus mucosa) and *ADH4* (esophagus muscularis, skeletal muscle) **(Methods).** Several other SNPs in strong LD with rs12639833 (rs2173201, rs2241894, rs3762896, rs6846835, rs10031168; R^2^ > 0.95, **S7 Table**) in YRI have extreme negative iHS scores, are eQTLs for increased *ADH1C* and *ADH4* expression, and were significantly associated with decreased risk for AD in African Americans. The selected alleles for three SNPs are associated with increased *ADH1C* expression in liver (rs3762896, rs6846835, rs10031168; personal communication, Y. Park, **S7 Table**). Taken collectively, these patterns suggest that (i) alcohol oxidation pathways broadly have been subject to recent positive selection in humans, (ii) that genes in this pathway have been repeatedly targeted, with multiple events segregating at these sites, (iii) the selective pressure appears to operate globally, at least in several populations from the major continental groups included in this study, and (iv) sweeping haplotypes at the *ADH* locus tag functional variation associated protection against alcohol dependence.

## DISCUSSION

We identified overlapping and shared signatures of positive selection across human populations, using a modified version of the statistic iHS that normalizes scores by local recombination rate. We observed more extreme iHS scores in sequencing data compared to SNP array genotype data, which could be a consequence of more rapid decay of homozygosity on unselected haplotypes due to the presence of rare variants. As expected, we found that closely related populations are more likely to share sweeping haplotype signatures, though we identified examples of sharing across genetically distant populations (e.g. African and non-African populations). These loci immediately raise questions of how these examples arose, whether by gene flow between continents after divergence or a common ancestral event. Though only a small amount of gene flow between African and non-African populations is thought to have occurred since their divergence, the introduction of an adaptively advantageous allele at very low frequency could lead to the signature we observe here. Future work modeling the potential scenarios leading to shared sweeps could help elucidate the evolutionary history of specific events.

One compelling example for recent positive selection involved the glycophorin A/B/E cluster in all four continental groups, with shared sweeps within African and South Asian groups, and additional independent sweeps in East Asian and European populations. To our knowledge, evidence of recent positive selection at this locus in these non-African populations has not been previously described. *GYPA* and *GYPB* encode glycophorin proteins, which reside on red blood cell membrane, and genetic variation in *GYPA* and *GYPB* determine individuals’ MNS blood group. *GYPE* is not known to encode functional protein, but may be a source of genetic variation for *GYPA* and *GYPB* in this region of frequent copy number variation and gene conversion [26,57,58]. Genetic variation in this region is associated with malarial resistance [27,30,59], and the signatures of positive and balancing selection that have been previously described at this locus may be due to the selective pressure of malaria on human populations. The malaria-resistant MNS blood type GP. Vw exists in Europe, possibly due to the endemicity of Malaria in Europe as recently as a few centuries ago [57]. Similarly, malaria-resistant MNS blood types are prevalent in East Asian populations with endemic *P. falciparum* malaria [57]. However, it is certainly conceivable that another infectious disease could take advantage of these cell surface proteins to invade erythrocytes, and be the selective agent in some (or all) of these populations. The frequency and diversity of apparent adaptive pressures at this locus underscores the role of selection on host-pathogen interactions over recent and longer evolutionary time-scales in modern humans, and the importance of this locus in particular in that process.

In a second case, our analysis indicated that oxidation of alcohol might have been subject to selective pressures more broadly across human populations than previously thought. At the alcohol dehydrogenase gene cluster in YRI, we identified a sweeping haplotype associated with decreased risk for alcohol dependence in African Americans, and increased expression of *ADH1C* and *ADH4* (in a multiethnic cohort of mostly European ancestry). This region has previously been shown to be under positive selection in East Asians and Europeans, but not African populations. Alcohol dehydrogenases oxidize ethanol to acetaldehyde, a process that is thought to occur primarily in the liver cells [63]. These data suggest a similar mechanism is at play in individuals of West African ancestry as in East Asians, where the selected allele increases *ADH* enzyme activity [55], resulting in an adverse physical response from alcohol consumption [60], and reduced risk for AD [53,56].

While our shared sweeps were not enriched for complex trait associations surveyed for a range of traits, we did find examples with a phenotype or variant that could be implicated. At the *DGKK* gene on chromosome X, we identified a sweep overlap across all five African populations that had a unique haplotype in three groups and a shared haplotype in two. In YRI, a risk variant for hypospadias, a birth defect of the urethra in boys, is in perfect LD with the sweep tag SNP. Multiple studies have found associations with hypospadias in the *DGKK* gene, and though none have been studied in a cohort of African ancestry it is clear that genetic variation in this gene plays a role in the development of the urethra in males. The clear adaptive potential of a variant that decreases risk for hypospadias, like the selected derived variant in YRI, provides a strong hypothesis for the phenotype under selection at this locus. In contrast, at the *MTHFR* locus in Europeans, the wide range of phenotypes associated with rs1801133 (a functional amino-acid changing mutation), make speculating on the endpoint phenotype under selection in Europeans difficult. In addition to higher homocysteine, lower folate, and lower vitamin B12 levels, the T allele has many reported associations including increased risk for multiple cancers [61,62], decreased risk for migraines [63], lower age of onset of schizophrenia [64], and increased risk for neural tube defects [65,66]. rs1801133 was also used as an instrument in a Mendelian randomization study that found evidence for a causal relationship between higher maternal homocysteine levels and lower offspring birthweight [67]. It is puzzling that the variant on the selected haplotype (T) is associated with a variety of maladaptive traits. If is truly underwent recent positive selection, it is possible that this variant is linked to another favorable allele, or this variant has some unknown highly favorable consequence that caused it to increase in frequency despite these associated detrimental phenotypes.

A final point of interest and a caveat to this work is the observed complexity of overlapping sweep regions. We frequently observed multiple sweeping haplotypes adjacent within a single population, and multiple overlaps across multiple populations in close proximity. Also common were independent sweeps across continental groups in the same location, a feature that could occur by chance or due to convergent evolution. We also found that the rate of sweep overlaps is not uniform across the genome, but in some locations overlaps cluster together, contributing to the complexity of the sweeps in those regions. These features made identifying the tag SNP for a sweep and calling sharing between sweep overlaps difficult in these regions. That said, we hope that our catalog of unusually long haplotypes shared across human populations will help to elucidate genes - and ultimately phenotypes - that are still evolving across the wide range of environments human have experienced in recent history.

## MATERIALS AND METHODS

### A correction to iHS adjusting for local, low recombination rates

While iHS was conceptualized for use in population data ascertained for common genetic variation [4], the empirical approach may not be calibrated on full-genome sequencing data where genetic variation across the allele frequency spectrum is more completely ascertained. To examine the properties of the score in more detail, we applied the iHS to population genetic data obtained from the Yoruba (YRI), CEPH (CEU), and Han Chinese (CHB) populations in the 1000 Genomes Project (1KG), Phase 3 (see below). First, we observed an excess of SNPs tagging strong iHS signals at lower derived allele frequencies (<20%, **S7A Fig**) in frequency range where iHS is not expected to have substantial power [4]. We observed a negative correlation between the number of populations in an overlap and the local recombination rate in any population (*e.g.,* Pearson’s correlation = 0.12, P = 9.9 × 10^−9^ in CHB). In addition, intervals tagged by these SNPs frequently overlapped across populations, particularly where the local recombination rate was also lower than the median rate genome-wide (*e.g.*, ρ = 1.4 × 10^−4^, P = 8.5 × 10^−10^ in CHB). After normalizing iHS by derived allele frequency and local recombination rate using a binning approach, summaries of the resulting score were much better calibrated to a mean of zero and unit variance (S8 Fig), substantially reducing though not abrogating the excess of high-scoring iHS values at low frequencies (S7B Fig). This normalization by local recombination rate also removed the association between low recombination rates and sweep overlaps across many populations. In all results described above, the iHS scores utilized this normalization scheme, treating autosomes separately from the X-chromosome (**S2 Table**).

### iHS scan

We downloaded phased genotype files for phase 3 of the 1000 Genomes Project from the 1KG FTP (http://ftp.1000genomes.ebi.ac.uk/vol1/ftp/release/20130502/). These data were converted to Beagle-formatted files, and filtered to include only biallelic SNVs (excluded indels) with a minor allele frequency (MAF) greater than 1%. A fine-scale recombination map was downloaded from the 1KG FTP (http://ftp.1000genomes.ebi.ac.uk/vol1/ftp/technical/working/20130507_omni_recombination_rates/), and scaled to units of ρ (=4N_e_r) for each population. Effective population size was estimated for each population by calculating nucleotide diversity (π) in a sliding window (100kb) across the genome, and estimating N_e_ from the median values π (N_e_ = π/(4*µ)). Ancestral alleles were identified using the human-chimp-macaque alignment from Ensembl (accessed from ftp://ftp.1000genomes.ebi.ac.uk/vol1/ftp/phase1/analysis_results/supporting/ancestral_alignments/). SNPs were filtered for only those where the ancestral allele was supported by both the chimp and macaque alignments.

Unstandardized iHS scores were calculated using WHAMM (v0.14a), using a modified version of iHS calculation code that increased speed of the calculation, and initially standardized by derived allele frequency as described in the original iHS paper [4], with 50 allele frequency bins. In the final standardization, we binned autosomal SNPs into 500 bins (50 allele frequency bins x 10 local recombination rate bins), or 150 bins for chromosome X (50 allele frequency bins x 3 local recombination rate bins). These standardization files are available in **S1 Table**.

Regions of the genome putatively undergoing recent hard sweeps - what we refer to in the main text as iHS intervals - were identified by counting the number of SNPs with |iHS| > 2 in 100kb windows (windows incrementing by one SNP, *i.e.*, overlapping windows). We took the union of the top 1% of windows, by the total number or by fraction of SNPs with |iHS| > 2 in the window, as our intervals. We performed this interval calling separately for each of the 20 populations included in this study. The SNP we use to label (*i.e.*, tag) each sweep interval was identified as the SNP with the most extreme iHS score, and the sweep frequency as the tag SNP derived allele frequency if the iHS score was less than zero, and ancestral allele frequency if iHS score was greater than zero. We limited our analyses of individual sweep loci to those with a tag SNP of MAF > 15%, to focus on signatures unlikely to have extreme iHS scores due to very low frequency.

### Sweep overlaps

To identify sweep overlaps, we compared the iHS intervals for each population and identified regions of the genome where two or more populations had a sweep interval. We calculated the fraction of sweep overlaps for each population pair as the mean of the fraction of sweep intervals in one population that overlap with a sweep interval in the second population (*i.e.* (fraction in pop A + fraction in pop B) / 2). We estimated F_ST_ for each pair of populations across all variants (n=2,627,240) in the 1000 Genomes VCF files on chromosome 2 using the Weir and Cockerham estimator implemented in VCFtools (v. 0.1.12b) [68]. Latitude and longitude for each population were estimated based upon the city listed by the 1KG project (*e.g.*, Tokyo for JPT) if the samples were collected at the site of ancestry, or by the approximate geographic center of the ancestral region if not sampled there (*e.g.*, Sri Lanka for STU). We performed stepwise forward regression in R (v. 3.3.1) on the fraction of sweep intervals overlapping between each pair of populations. Possible predictor variables were difference in latitude, difference in longitude, straight-line geographic distance, F_ST_, and the continental group labels (*e.g.*, EUR vs. EUR = both populations within Europe or EUR vs. SAS = one European and one South Asian population).

### Rates across the genome

To assess the rate of sweep intervals across the genome, we subdivided the genome into ten megabase non-overlapping windows (n=297 in total) and counted the number of sweep intervals for each individual population, and the number of overlaps across 2 or more populations, in each window. To ensure the sweep intervals called for each population were independent, we merged adjacent sweep intervals into one interval if their tag SNPs were in modest LD or greater (R2 > 0.4). We used all Ensemble HG19 gene annotations (from http://genome.ucsc.edu/cgi-bin/hgTables), merged into non-overlapping intervals with BEDTools v2.19.1 [69]. If a sweep interval or overlap spanned two windows, we counted it once in the window with more than half of its physical distance. We fit mixtures of independent Poisson distributions to the data by minimizing the negative log likelihood with the non-linear minimizer function (nlm) in R v. 3.3.1 [70]. We compared mixture models by calculating the Bayesian information criterion and performing a likelihood ratio test.

### Identifying sweeping haplotypes with fastPHASE

For each sweep overlap, we identified the physical region spanning all tag SNPs, and an additional 5kb to either side. We ran fastPHASE on this region, using the -u option to identify each 1KG population as a subpopulation, -B to indicate known haplotypes, and -Pzp to output cluster probabilities for each individual at each SNP. We tested a range of values of K (number of haplotype clusters) and T (number of random EM algorithm starts) on a subset of sweep overlaps, and found broadly similar results across the range (data not shown). We used K = 10 clusters and T = 10 for all overlaps in the final analysis. From the output cluster probabilities, we identified the sequence of haplotype clusters for each SNP position in each individual as the most likely haplotype cluster at each SNP. We then identified the haplotype cluster sequences of all chromosomes carrying the selected tag allele, and the most common of those to be the reference sweeping haplotype sequence.

To identify if a pair of populations as “shared”, we required an identical reference haplotype sequence to span the selected tag allele in both populations. To form shared clusters, we grouped together all populations that were called as shared with at least one other population. To calculate the null rate of haplotype sharing across population pairs, we selected random regions of the genome of the same size and distance to genes as our observed sweep overlap regions. For each sweep overlap, we identified 10 matched windows, for a total of 30,450 regions across the genome (ranging from 153-2588 random overlaps per population pair). We identified tag SNPs for each population in the random regions matching the distance from the other populations’ tag SNPs and derived allele frequency (within 5%) of the observed overlap. We then ran fastPHASE on the randomly selected regions and performed the shared haplotype-calling procedure as for observed overlap windows described above. To compare the observed fraction of overlaps called as shared to the null haplotype sharing for each pair of populations, we performed 1000 bootstraps by sampling with replacement the number of observed overlaps from the null. Population pairs where the shared sweep fraction of observed overlaps was higher than the shared fraction of random overlaps for all 1000 samples are marked with an asterisk in Fig 3.

### Enrichment/GTEx

To connect shared sweeps to potential causal genes, we utilized the GTEx v6 eQTL dataset downloaded from the GTEx portal (http://www.gtexportal.org/) [13]. For each population’s tag SNPs, we identified LD proxies (R^2^ ≥ 0.9, calculated in the same population) within 1 Mb of the sweep interval, and intersected these SNPs with all significant GTEx eQTLs from all tissue types. eQTLs in the GTEx V6p data set were identified using a cohort of mostly white individuals (84.3%), with a smaller fraction of African Americans (13.7%). For sweep overlaps that were called as shared, we identified a shared SNP set as the intersection of LD proxy sets for all populations in a shared group. We created a gene list of all genes with eQTLs from any tissue that intersected with shared SNP sets, excluding HLA genes. We chose to exclude HLA genes, owing to its genomic complexity and its enrichement for signatures of recent positive selection. To test for enrichment of this gene set with biological pathways, we used over-representation analysis of all pathway databases in ConsensusPathDB (http://cpdb.molgen.mpg.de/) [48] with the background set of all genes.

### Intersection of sweeps with Neandertal haplotypes, Neandertal PheWAS, and GWAS SNPs

We downloaded the Neandertal haplotype calls reported in [19] from http://akeylab.gs.washington.edu/vernot_et_al_2016_release_data/ (no X chromosome data available). To calculate LD between introgressed haplotypes and sweep tag SNPs, we pooled overlapping haplotypes across individuals and created a genotype of 0 or 1 based on presence/absence of the overlapping introgressed haplotype in each individual. We then calculated LD between this presence/absence genotype and the tag SNPs within 1 Mb of the introgressed haplotype separately for each population. We considered haplotypes with R^2^ > 0.6 with sweep tag SNPs as candidates for adaptive introgression. To examine potential enrichment of introgressed haplotypes in LD with sweep tag SNPs, we compared the fraction of introgressed haplotypes in LD with sweep tag SNPs to the distribution of all SNPs within 1 Mb of tag SNPs with R^2^ > 0.6. We downloaded the Neandertal PheWAS data at https://phewascatalog.org/neanderthal [45], and intersected all reported associations with variants in strong LD (R^2^ ≥ 0.9) with each sweep tag SNP in each population.

We downloaded the GWAS catalog from https://www.ebi.ac.uk/gwason10/12/16. We identified all genome-wide significant associations (P < 5 × 10^−8^) in strong LD (R^2^ ≥ 0.9) with each sweep tag SNP in each population. To test for enrichment of GWAS variants generally and of specific phenotype classes, we performed permutation tests with random SNP sets from the HapMap3 variant set (from ftp://ftp.ncbi.nlm.nih.gov/hapmap/phase_3/) matched for allele frequency and distance to gene with the GWAS variants of interest. We then compared the empirical distribution of intersection of these matched SNP sets with the sweep tag SNPs and proxies to the number of observed GWAS intersections. To control for potentially linked GWAS variants, we simply counted the number of sweeps in each population that intersected a GWAS or control set variant.

### Indels and annotations

Indels were not included in our iHS scan, but could be the causal variant on a sweeping haplotype. To identify candidates for causal indels, we calculated LD with sweep tag SNPs for all indels in the 1000 Genomes phase 3 VCF files within 1 Mb of the sweep interval in each population. To identify potential functional coding variants among indels and SNPs on sweeping haplotypes, we used ANNOVAR to annotate coding variation [71].

## ACKNOWLEDGEMENTS

This work was supported through grants from the National Institutes of Health (NIDDK R01DK101478) and a fellowship from the Alfred P. Sloan Foundation (BR2012-087) to BFV. KEJ was supported in part by National Institutes of Health T32GM008216.

### SUPPLEMENTARY FIGURES

**S1 Fig.**
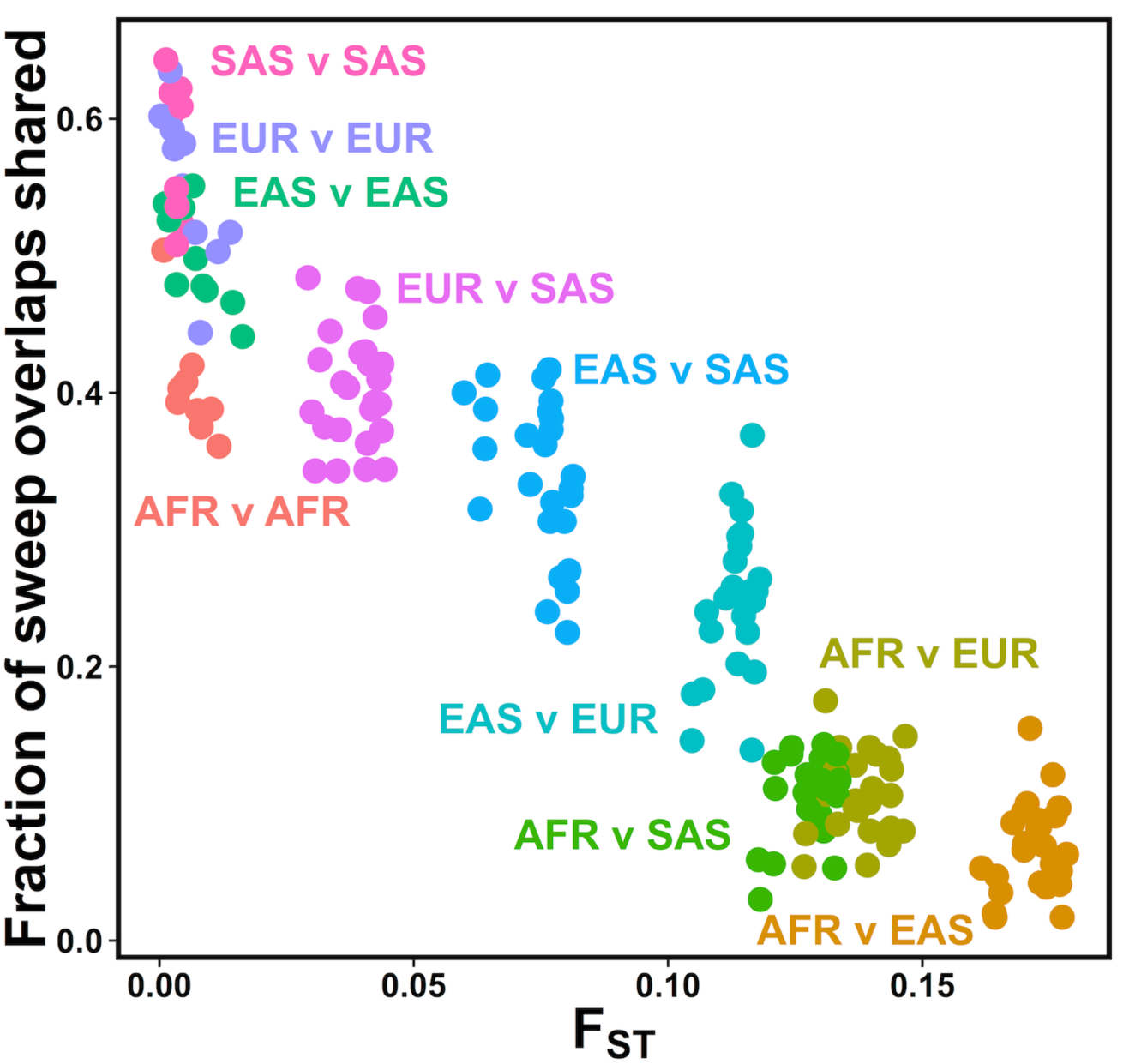
Closely related populations have shared sweeps more frequently. For each population pair, the fraction of sweep overlaps that are shared is plotted against pairwise estimated F_ST_. Each population pair (dots) are colored by their continental groupings (e.g. EUR v SAS = one European population *vs*. one South Asian population).

**S2 Fig.**
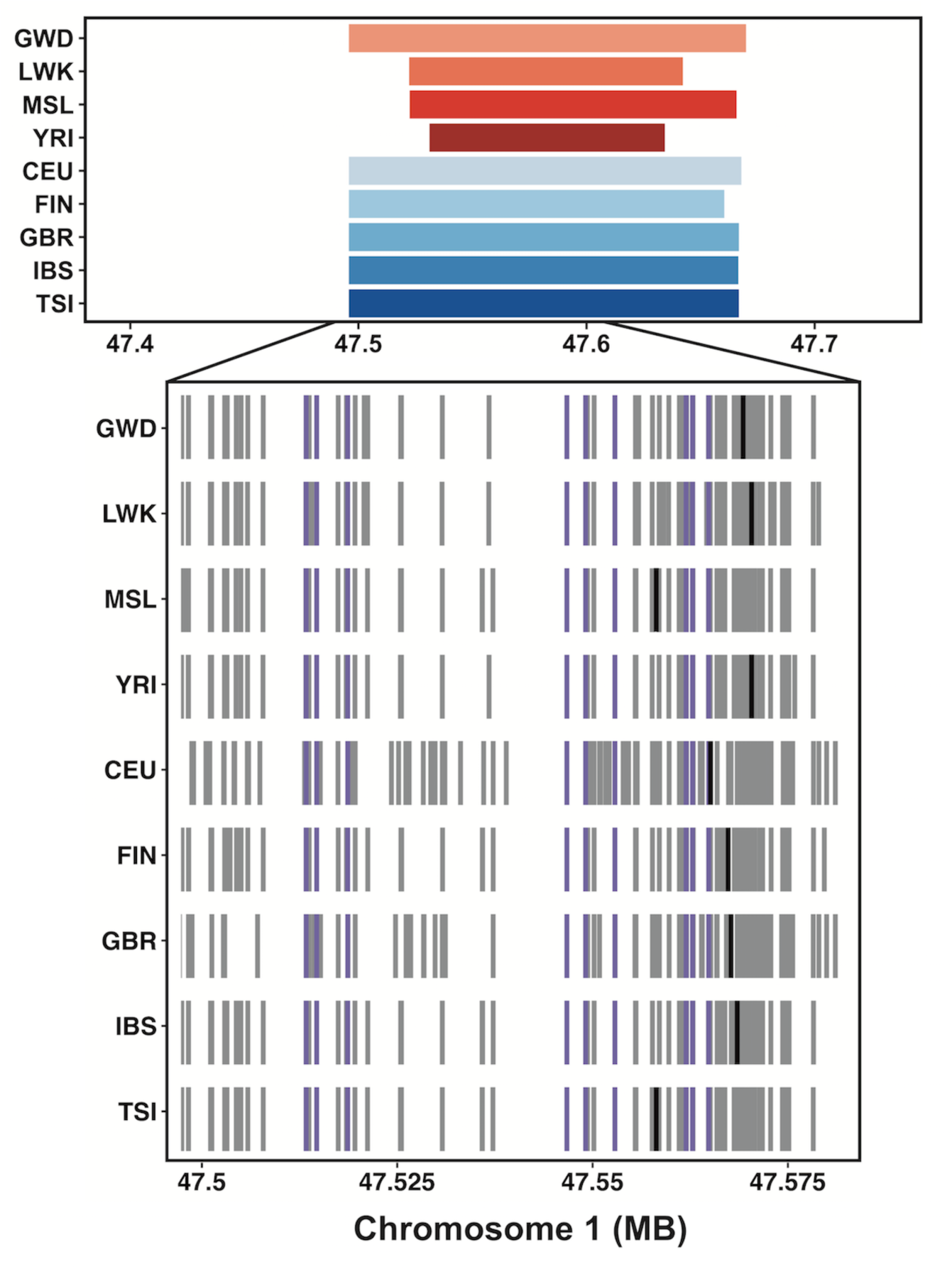
Signatures of positive selection at the cytochrome P450 locus on chromosome 1. We observed 9 populations from the European and African continental groups with a shared sweep at this locus. The top panel shows the genomic positions of each populations’ iHS interval. The bottom panel shows the variants on the most common sweeping haplotype in each population, with gray ticks indicating the presence of derived variants. The location of the highest-scoring iHS SNP in each population is shown with a black tick. Nine SNPs in LD with the shared haplotype of these nine populations (D’ = 1) were eQTLS for *CYP4X1*, *CYP4Z1*, and *CYP4A22-AS1* in testis, and their positions are shown in purple.

**S3 Fig.**
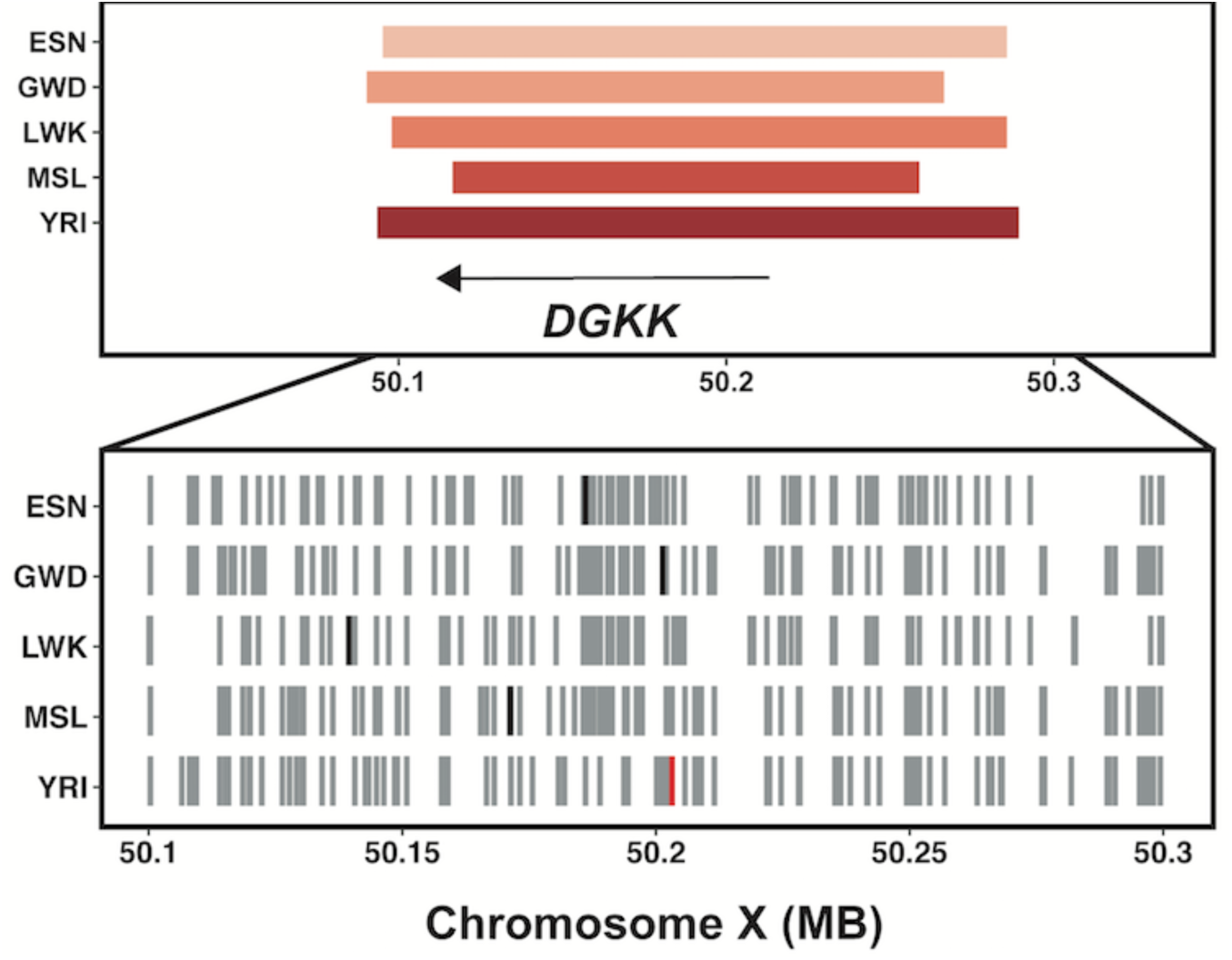
Signatures of positive selection at *DGKK* on chromosome X. The top panel shows the iHS intervals for five African populations with sweeps at this locus, and the position of the *DGKK* gene. The bottom panel shows the variants on the most common sweeping haplotype in each population, with gray tick marks indicating derived alleles, and black ticks marking the SNP with the most extreme iHS score in each population. The highest scoring SNP in YRI (red tick) was rs4554617, a SNP associated with hypospadias in GWAS. MSL and LWK have a shared sweep in this overlap.

**S4 Fig.**
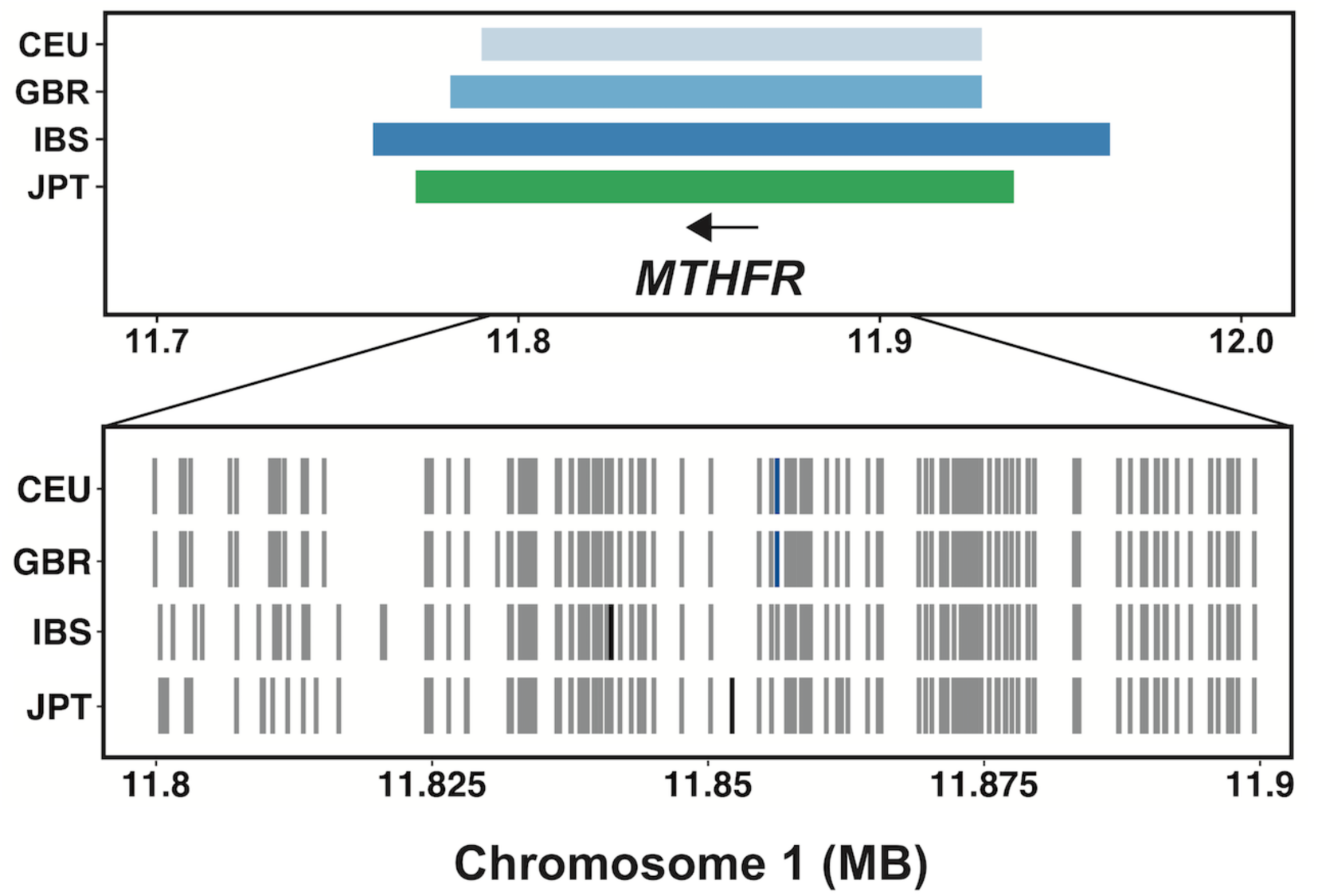
Signatures of positive selection at *MTHFR* on chromosome 1. The top panel shows the positions of iHS intervals for three European populations and one East Asian population with sweeps at this locus, and the position of the *MTHFR* gene. rs1801133, a coding variant in *MTHFR*. The bottom panel shows variants on the most common sweeping haplotype in each population, with derived alleles shown in gray and the SNP with the most extreme iHS value marked in black or blue. The highest-scoring iHS SNP in CEU and GBR, rs1801133 (position marked with blue ticks), is a coding variant in *MTHFR* associated with numerous traits.

**S5 Fig.**
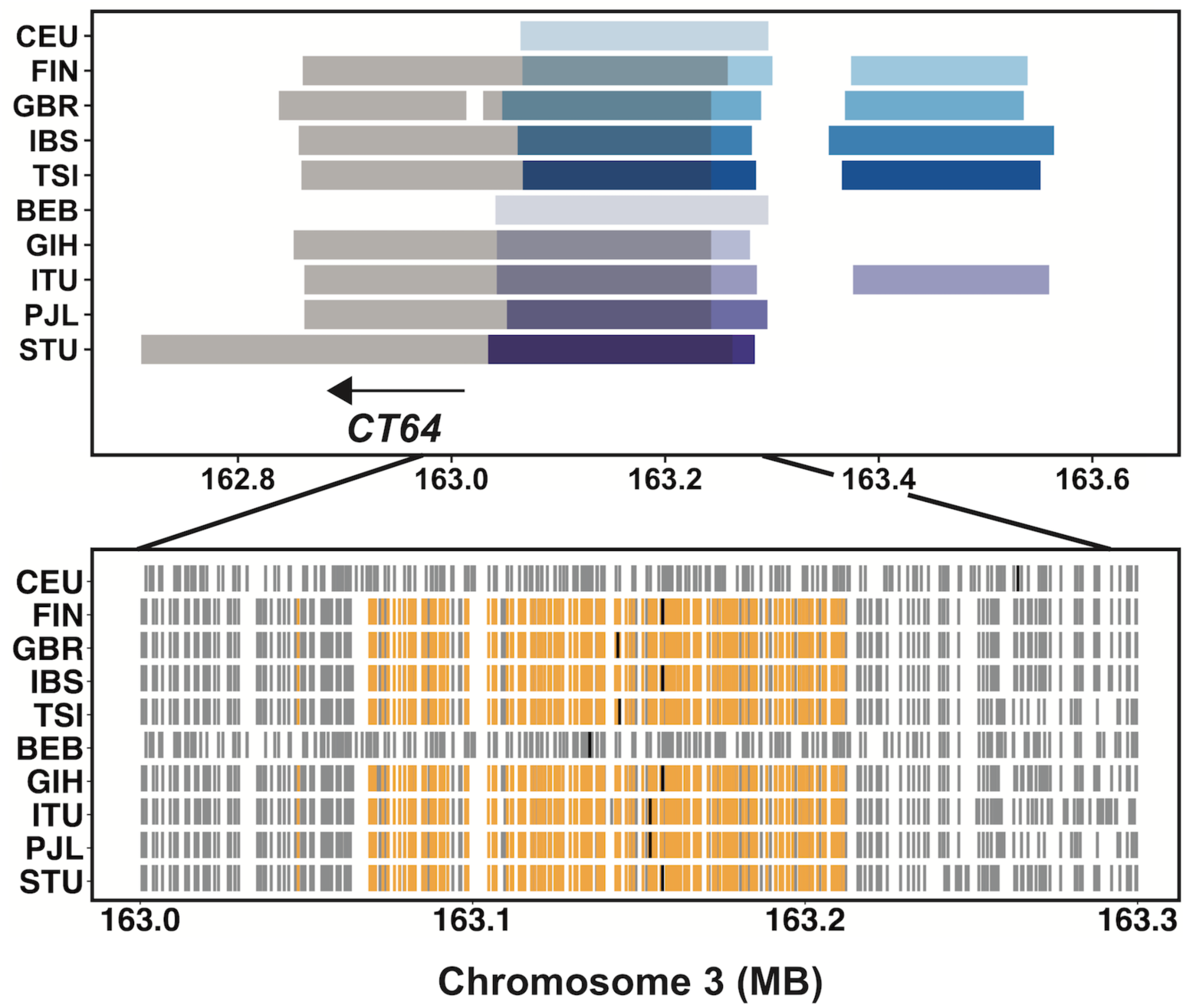
Signature of positive selection and introgressed Neandertal haplotypes at a testis-expressed non-coding RNA. The top panel shows the sweep intervals for European (blue) and South Asian (purple) populations, the positions of linked (R^2^ > 0.8) introgressed Neandertal haplotypes (transparent gray), and the position of non-coding RNA *CT64*. The bottom panel shows the most common sweeping haplotype in each population, with gray ticks indicating derived variants. The position of the SNP with the most extreme iHS score in each population is shown with a black tick. The positions of eQTLs for the non-coding RNA *CT64* from testis in LD (R^2^ > 0.9) with the shared sweeping haplotype in eight populations are shown with orange ticks.

**S6 Fig.**
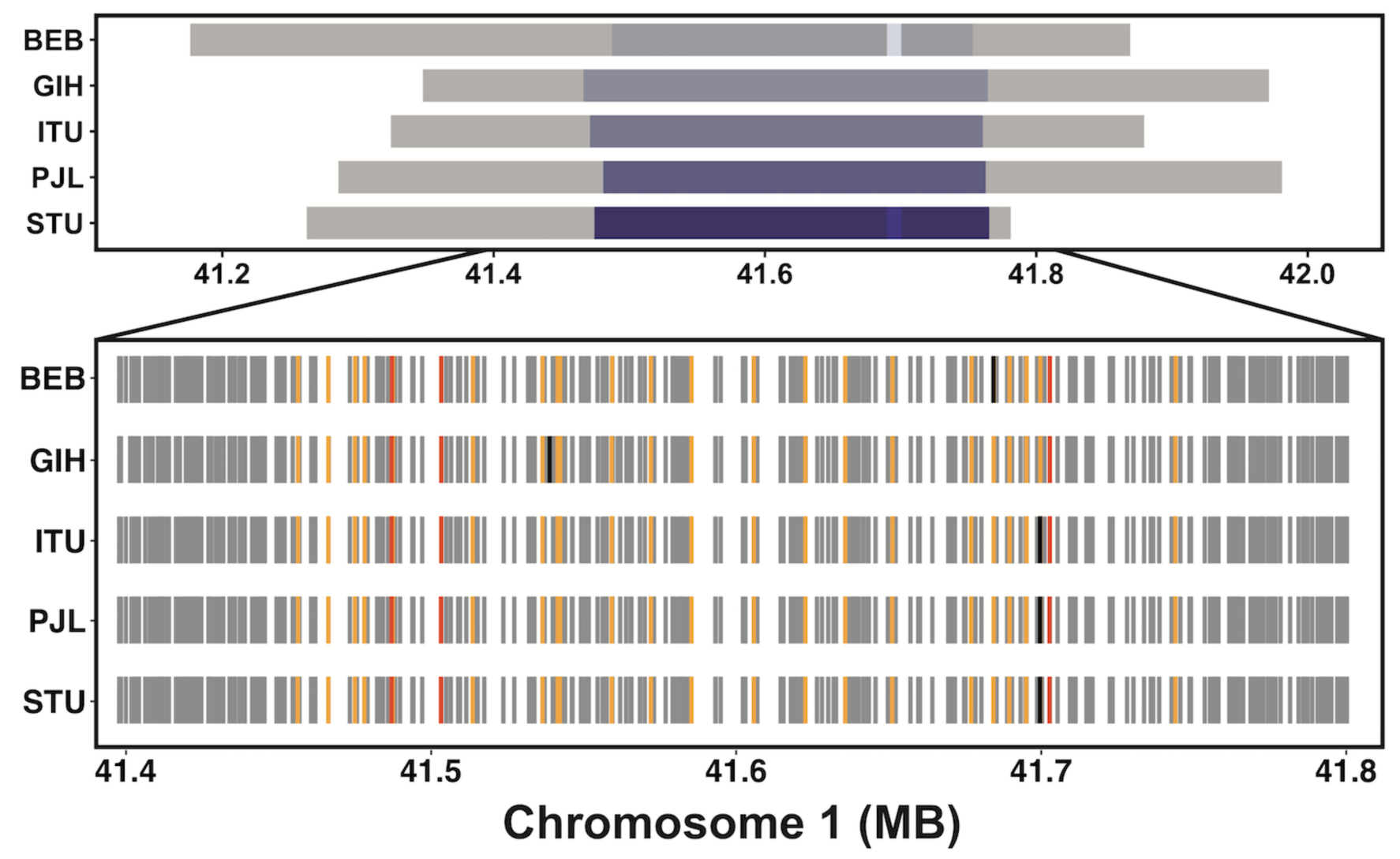
Signatures of positive selection and introgressed Neandertal haplotypes at chromosome 1: 41 MB. The top panel shows the positions of iHS intervals for five South Asian populations (purple), and the positions of linked (R^2^ > 0.25) introgressed Neandertal haplotypes (transparent gray). The bottom panel shows variants on the most common sweeping haplotypes in each population, with gray ticks indicating derived alleles. The SNP with the most extreme iHS score in each population is shown with a black tick. Also shown are the positions of pheWAS SNPs (dark orange) and 22 eQTLs for five genes (light orange) linked to the shared sweeping haplotype (R^2^ > 0.9). The five genes and their eQTL tissues are: *RP11-399E6* = lung, *SLFNL1* = brain cerebellum, *SCMH1* = esophagus muscularis, *FOXO6* = brain cerebellum, *CTPS1* = testis.

**S7 Fig.**
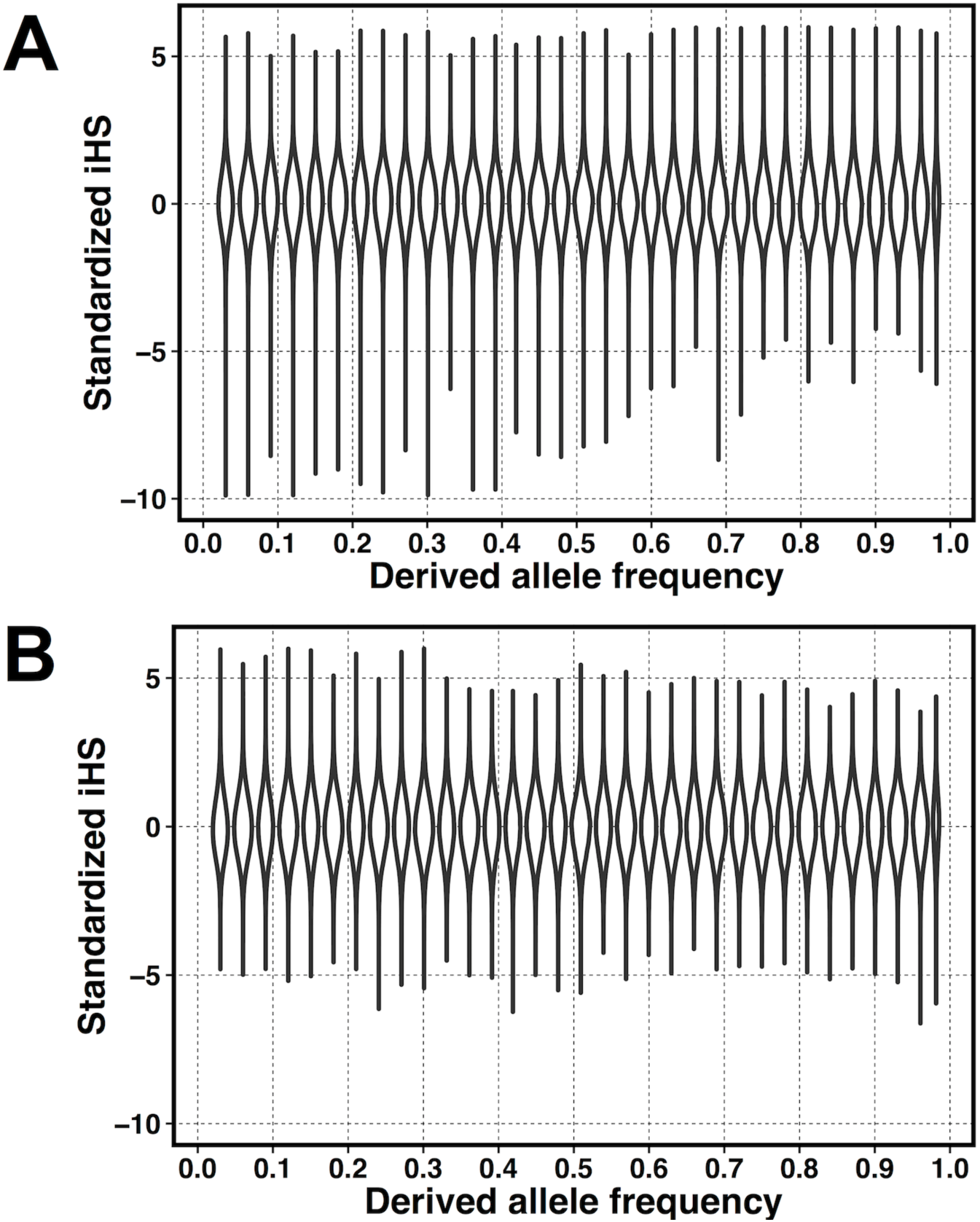
Standardized iHS scores and derived allele frequencies. The distribution of standardized iHS scores in YRI as a function of derived allele frequency before (A) and after (B) recalibration for local recombination rate. Each violin plot represents a bin of SNPs within a 3% range of derived allele frequencies.

**S8 Fig.**
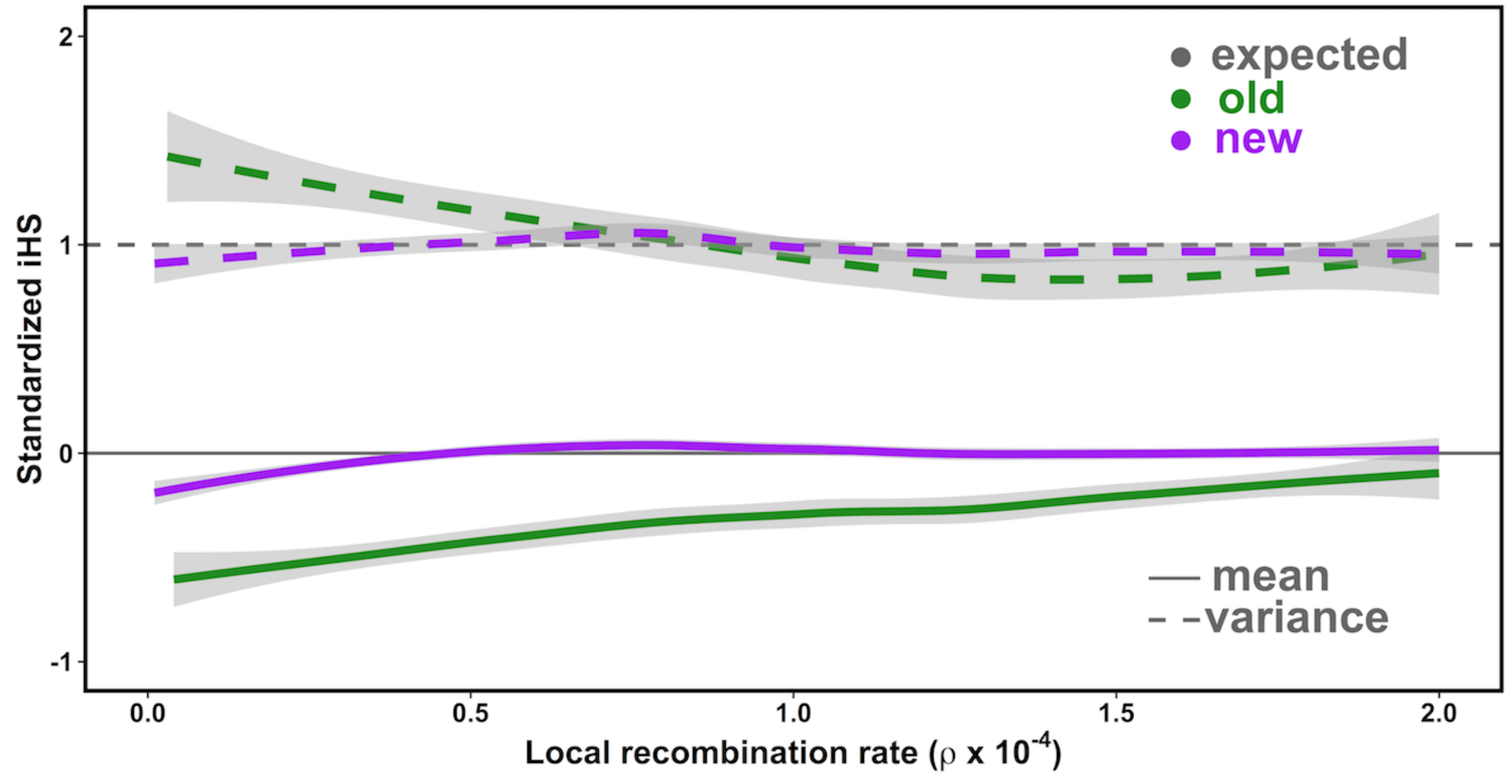
Normalizing iHS by local recombination rate. The mean (solid line) and variance (dashed lined) of iHS scores as a function of local recombination rate. iHS scores normalized by derived allele frequency are shown in green; normalization by both derived allele frequency and local recombination rate shown in purple. The gray lines represent a mean (= 0) and variance (= 1) for comparison.

### SUPPLEMENTARY TABLES

**S1 Table. 1000 Genomes population codes.**

**S2 Table. iHS standardization tables.** Each population has two tables, one for autosomes and one for the X chromosome.

**S3 Table. iHS intervals and tag SNPs for all populations.**

**S4 Table. iHS interval count and estimated effective population sizes.**

**S5 Table. Sweep intervals shared across populations.**

**S6 Table. Sweep sharing for each population pair.** For each population pair, the fraction of sweep overlap sharing, along with the background sharing rate and bootstrapped 99% confidence interval.

**S7 Table. Alcohol dependence GWAS SNPs on the sweeping haplotype at the ADH locus in YRI.** For each SNP, the iHS scores, derived & ancestral alleles, association with alcohol
dependence (in Af. Am.) [54], eQTL beta and p-values [13], R2 with lead GWAS SNP (in YRI
& ASW).

**S8 Table. GWAS SNPs linked to iHS signatures.** All pairs of SNPs with R2>0.9 between a
populations' iHS tag SNP and a GWAS SNP are listed. riskSel: is allele on selected haplotype (derived for negative iHS score, ancestral for positive iHS score) the GWAS risk allele?

**S9 Table. Introgressed Neandertal haplotypes linked to iHS tag SNPs** (R2>0.6).

The iHS code used in this paper can be found at https://github.com/bvoight/iHS_calc
Standardized iHS scores for every population and SNP included in this study can be found at coruscant.itmat.upenn.edu

